# Lateralized role of prefrontal cortex in guiding orienting behavior

**DOI:** 10.1101/2020.04.12.038356

**Authors:** Ali Mohebi, Karim G. Oweiss

**Affiliations:** Department of Neurology, University of California San Francisco, San Francisco, CA; Electrical & Computer Engineering, Biomedical Engineering, Herbert Wertheim College of Engineering; Department of Neuroscience, the McKnight Brain Institute; Department of Neurology, the Norman Fixel Institute for Neurological Disorders, University of Florida, Gainesville, FL

## Abstract

Orienting movements are essential to sensory-guided reward-seeking behaviors. Prefrontal cortex (PFC) is believed to exert top-down control over a range of goal-directed behaviors and is hypothesized to bias sensory-guided movements. However, the nature of PFC involvement in controlling sensory-guided orienting behaviors has remained largely unknown. Here, we trained rats on a delayed two-alternative forced-choice task requiring them to hold an orienting decision in working memory before execution is cued. Medial PFC (mPFC) Inactivation using either Muscimol or optogenetics impaired choice behavior. However, optogenetic impairment depended on the specific trial epoch during which inactivation took place. In particular, we found a lateralized role for mPFC during the presentation of instruction cues but this role became bilateral when inactivation occurred later in the delay period. Electrophysiological recording of multiple single-unit activity further provided evidence that this lateralized selectivity is cell-type specific. Our results suggest a previously unknown role of mPFC in mediating sensory-guided representation of orienting behavior and a potentially distinct cell-type specific role in shaping such representation across time.

## INTRODUCTION

Movement is pivotal to reward-seeking behaviors. To obtain rewards, animals have to continuously move to sense their environment. Since movement is costly, both in terms of energy expenditure and time, significant brain resources have been dedicated to optimizing goal-directed movements. The prefrontal cortex (PFC), in particular, has been implicated in guiding goal-directed actions (Miller and Cohen, 2001; Quintana and Fuster, 1999; Szczepanski and Knight, 2014) among other cognitive functions that serve that purpose such as attention modulation, working memory, rule learning and task switching (Donahue and Lee, 2015; Duan et al., 2015; Fujisawa et al., 2008; Greene et al., 2001; Romo and de Lafuente, 2013). Receiving sensory input from the mediodorsal nucleus of the thalamus and projecting to the premotor cortex, PFC is believed to gate in relevant information for sensorimotor integration (Euston et al., 2012; Gabbott et al., 2005; Narayanan and Laubach, 2006). It has been proposed that PFC influences motor goal representation by encoding rules and strategies that govern reward seeking actions (Gisquet-Verrier and Delatour, 2006; Karlsson et al., 2012; Powell and Redish, 2016; Rich and Shapiro, 2009). However, the extent to which this top-down influence affect more basic actions such as orienting the head, eyes and body has remained largely unknown.

Orienting behavior is a primal stereotyped movement essential to directing attention towards salient sensory input, which in turn facilitates foraging, navigation among other behaviors. Numerous cortical and subcortical structures involved in orienting behavior have been identified in rodents. In particular, lesion and electrophysiological recording studies have established the superior colliculus (SC) and frontal orienting field (FOF) as critical for orienting execution and planning respectively (Duan et al., 2015; Erlich et al., 2011; Kopec et al., 2015). However, the role of medial PFC in guiding orienting behaviors in rodents has remained poorly understood.

Here, we used a delayed two-alternative forced-choice task (2-AFC) to study the role of rat mPFC in sensory-guided orienting behaviors. In particular, we first show that prolonged inactivation of mPFC leads to impaired performance. This inactivation affected both choice accuracy and reaction time. We then demonstrate that this impairment is inactivation time-dependent: inactivating mPFC during cue presentation only affects contralateral choices whereas inactivation during motor planning affects both choices. Finally, using multielectrode recording we show how goal formation evolves in the dynamics of mPFC activity.

## RESULTS

### Two-alternative forced-choice task

We trained a total of n=19 rats to perform a two-alternative, forced-choice task. Details of behavioral training are described elsewhere (Mohebi and Oweiss, 2014). Briefly, adult Sprague-Dawley rats (Charles-River) were food deprived to their 85% ad libitum weight and placed in acoustically isolated training boxes (Coulbourn instruments, PA). The box comprised an operant conditioning chamber with three nose poke holes (a fixation hole in the center and two target holes on the sides) and a reward delivery trough on the opposite side of the nose poke holes as shown in Figure 1a. Subjects self-initiate a trial by placing their snout inside the fixation hole. Following a 1s fixation period, a brief (500 ms) single-frequency tone (60 dB) was played. Pitch of the tone instructed the rat to make a choice to their rat or left. The instruction cue was followed by a delay period of variable length (1-1.5s), which was then immediately followed by a Go cue (white noise tone). Premature retractions terminated the trial. Correct choices were rewarded by delivering a 45mg food pellet in the trough. Rats adapt their choice depending on the instruction cue. Choice behavior is captured by a sigmoid function on both individual and group level behavior (Fig. 1b, n=13 sessions, β = 0.62 ± 0.23).

**Fig. 1.**
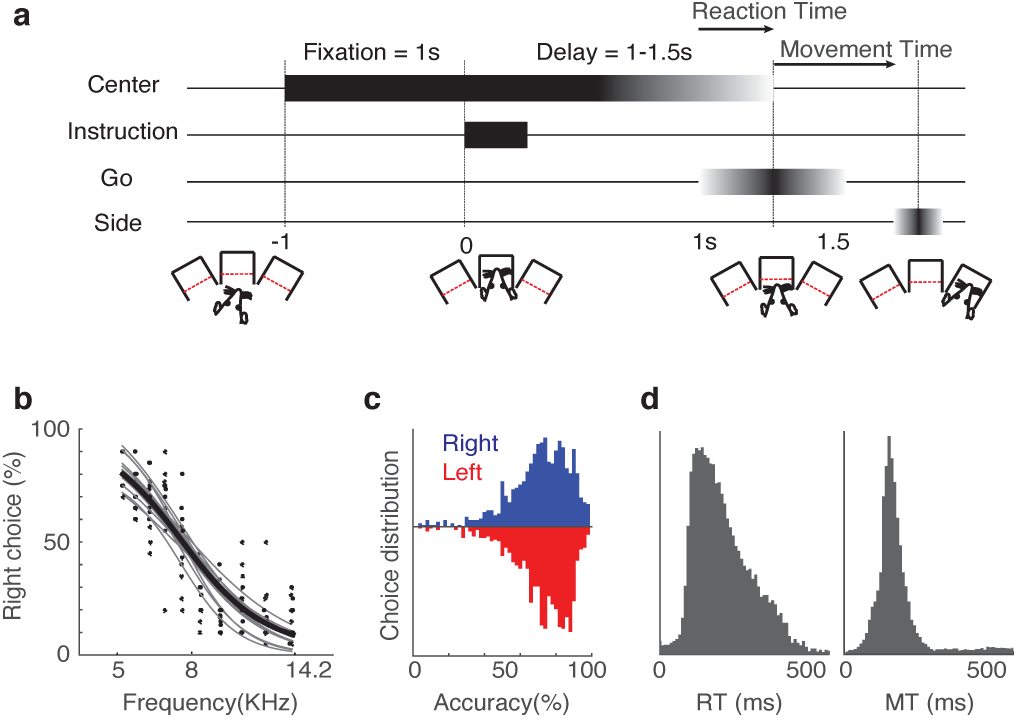
Rats perform a delayed auditory two-alternative choice task. **a.** Schematic showing the trial timeline. Rats poke in a center hole and wait for the auditory instruction cue. The pitch of the tone instructs the rewarded choice. Rats have to hold in the center port until the go cue. **b.** Choice performance showing the distribution of choice accuracy over frequency. Data is collected from 13 sessions. **c.** Choice performance under the two-tone version of the task (5, 14.1 KHz). Data collated from 1039 sessions showing significant performance above chance level (p<10^−6^, binomial test) **d.** Distribution of reaction time (RT) and movement time (MT).

We only used the two extremes of the stimulus spectrum (5KHz and 14.2 KHz associated with right and left choice, respectively) throughout the rest of the study. Rats reached > 85% performance after an average of 30±8 sessions (Fig. 1c, n=1039 session, p<10^−6^, binomial test).

### Unilateral inactivation of mPFC impairs choice behavior

We locally infused muscimol -a GABA_A_ agonist - to reversibly inactivate the dorsal prelimbic cortex (Baker and Ragozzino, 2014; Willcocks and McNally, 2013). Microinjection was delivered unilaterally through a chronically implanted cannula (Fig. 2a) and drug infusion sessions were interleaved with sham injections (ACSF). We found that subjects showed selective impaired choice following drug injection. In particular, choices ipsilateral to injection were as accurate as the control group; however contralateral choices were selected at significantly lower accuracy (Fig. 2b, Mann-Whitney U test, p<10^−3^). In addition to this ipsilateral choice bias, reaction times were significantly slower under the muscimol administration (Fig. 2c, paired KS-test,p<10^−5^). Unlike choice behavior, however, slowing down of reaction times was not selective and manifested in both contralateral and ipsilateral movements.

**Fig. 2.**
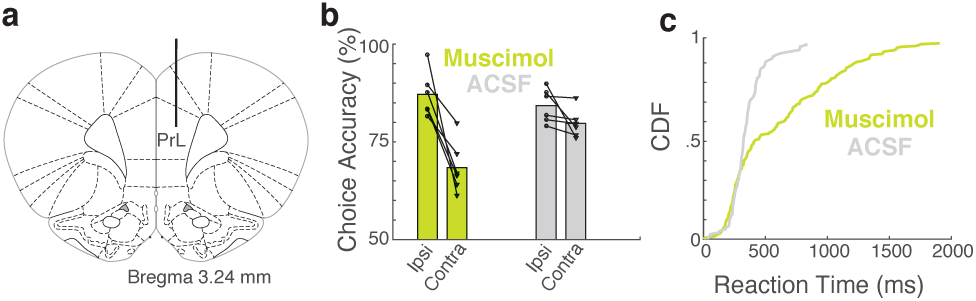
Reversible inactivation of the prelimbic cortex impairs choice behavior. **a.** Schematic showing infusion cannula placement targeting the dorsal prelimbic cortex for muscimol delivery. **b.** Choice accuracy is significantly reduced for choices contralateral to muscimol infusion side (Mann-Whitney U test, p<10^−3^) and not ACSF. **c.** Muscimol infusion slowed down reaction times (RT) compared to ACSF injections. The graph shows the cumulative distribution function (CDF) of reaction times. (paired KS-test, p<10^−5^).

### Optogenetic dissection of mPFC activity reveals separable states

Choice impairment and slowing down of movements following muscimol injection suggests a role for mPFC neurons in guiding orienting behavior during the task, similar to previous reports (Erlich et al., 2011). However, pharmacological inactivation lacks the temporal precision to dissect the exact dynamics of mPFC during different trial epochs. We then resorted to an optogenetic approach to perturb neural activity during specific task epochs. We expressed the inhibitory opsin ArchT3.0 in excitatory cells of PFC under the CaMKII promoter and used green laser (532nm, 10mW) to inactivate pyramidal cells in mPFC during instruction, delay and action execution epochs.

First, we sought to recapitulate the inactivation effects observed during the muscimol inactivation experiments. We suppressed the activity of mPFC pyramidal in a fraction of trials (25% selected in random order) for the entire trial period (Fig. 3b). This led to contralateral choice impairment and non-selective slowing down of reaction times (Fig3c-e), similar to our pharmacological inactivation results. Using blue light as a control (473nm, 10mW) did not affect the choice or reaction time (Fig. 3c).

**Fig. 3.**
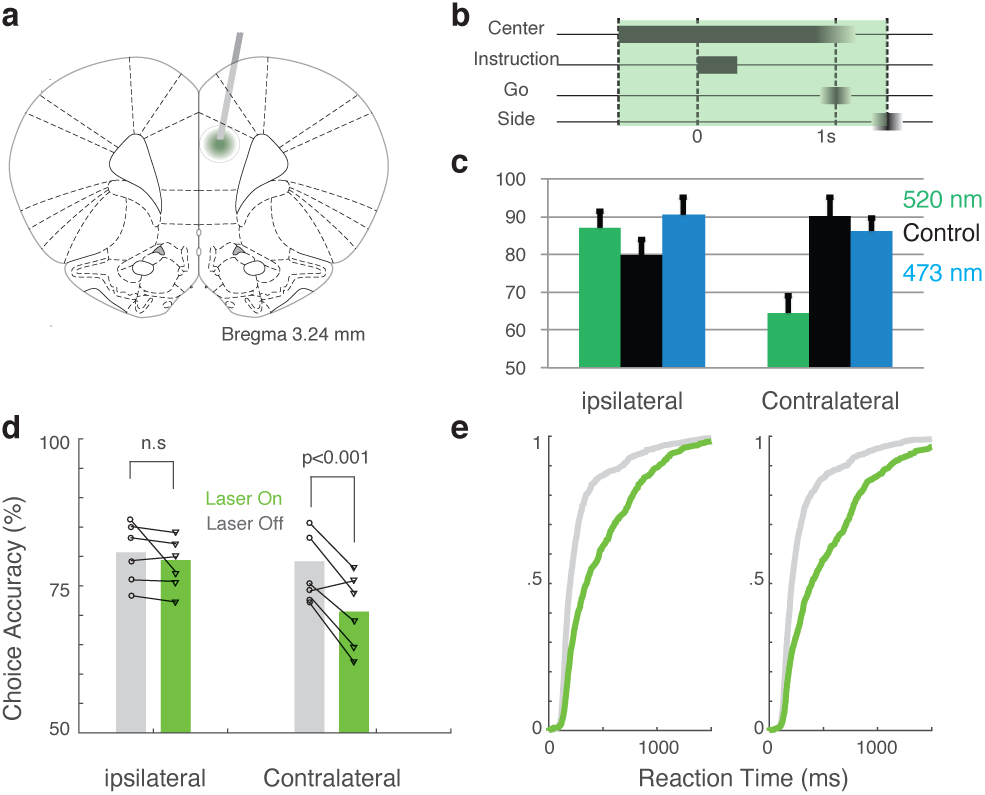
Optogenetic inhibition of the prelimbic cortex reduces choice accuracy. **a.** Schematic showing injection target and optical fiber placement. **b.** A 10Mw laser pulse was delivered throughout the trial, starting at the fixation until side choice. **c**,**d.** ArChT activation under green laser pulse (520nm, 10mW) significantly reduced choice accuracy for contralateral trials (Mann-Whitney U test, p<10^−3^). Blue light (473nm, 10mW) trials accuracy, however, was not significantly different from control trials **e.** Reaction times are non-selectively slowed down in laser trials (KS-test, p<10^−4^).

Next we limited the inactivating laser pulse to specific epochs of the task. Unilateral, optogenetically-induced suppression (500ms pulses) limited to the instruction period impaired choice performance and slowed down reaction time, but only for contralateral choice trials (Fig. 4a-c). Inactivation during delay and reaction period, on the other hand, equally impaired choice accuracy and slowed down reaction times for both ipsilateral and contralateral choices (Fig. 4d-i).

**Fig. 4.**
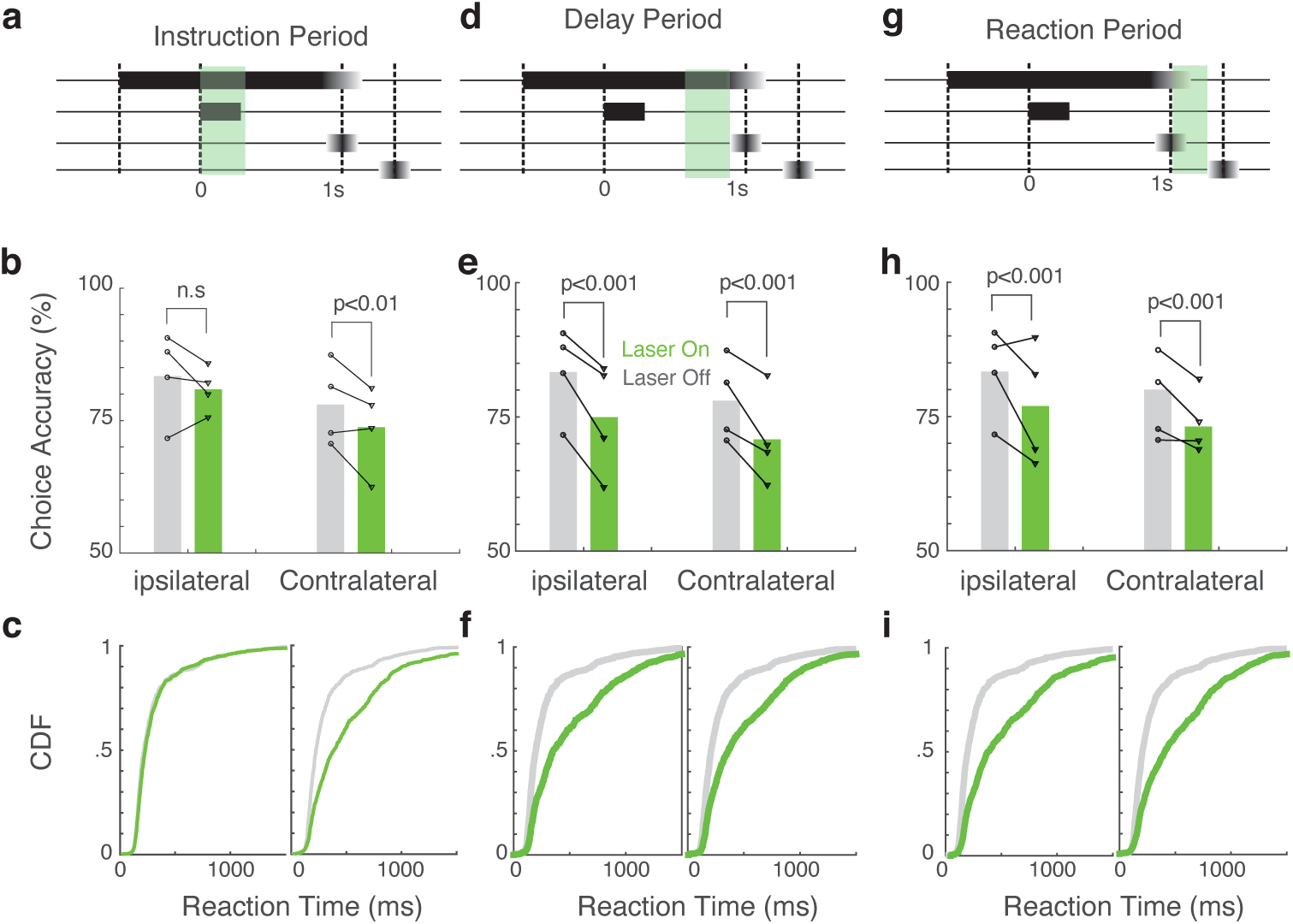
Brief inactivation of prelimbic cortex during critical epochs results in lateralization of upcoming choice. **a.** Schematic showing the time course of optogenetic suppression of activity during the instruction period. **b.** Choice accuracy comparing laser trials to non-laser trials for contralateral and ipsilateral choices. **c.** Cumulative distribution of reaction times for laser and non-laser trials for ipsilateral and contralateral choices **d-f.** Similar to a-c but for optogenetic suppression during the late delay period (.5s before Go cue). **g-i.** Similar to a-c but for optogenetic suppression delivered during the choice period (at the Go cue onset until the side choice)

### Lateralized representation of choice among mPFC cells

We observed that suppression of mPFC during the instruction epoch caused selective impairment in contralateral choices whereas similar inactivation during delay and reaction epochs caused a non-selective impairment. Given this perplexing observation, we hypothesized that during the instruction period, only contralateral orienting information is represented by mPFC activity and ipsilateral representations evolve at later timepoints. To test this hypothesis, we used chronically implanted microwire arrays (Tucker Davis Technologies, Alachua, FL) to record from many single neurons in prefrontal cortex simultaneously during the task, sampling a broad range of layer V dorsal prelimbic cortex along the rostro-caudal axis. We recorded wideband signals and then isolated single units using offline analysis (Fig. 5a).

**Fig. 5.**
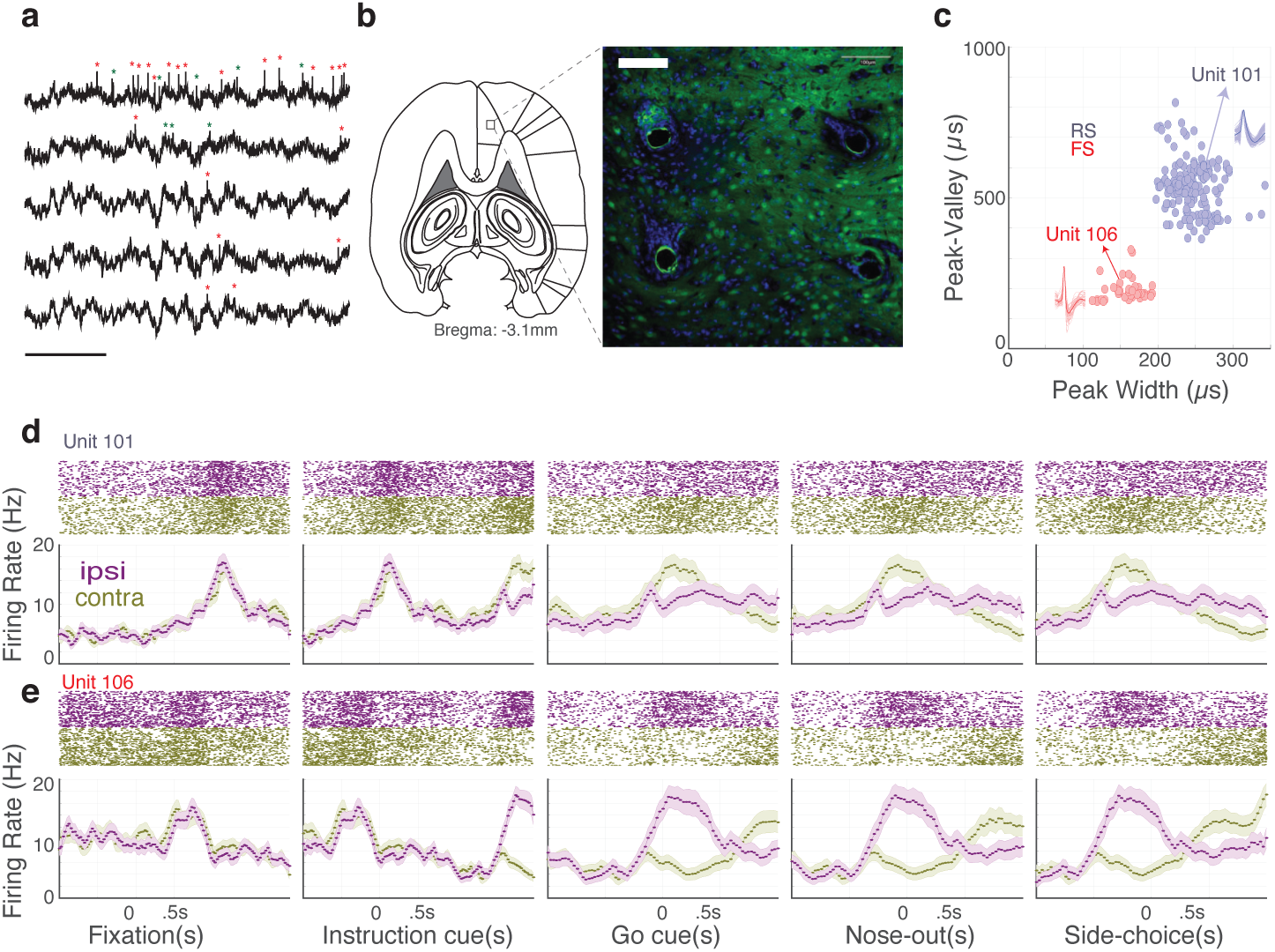
Single unit recording in mPFC reveals heterogeneous response patterns among individual units. **a.** Representative trace of wideband recording across multiple channels revealing slow waves as well as multiple single units (Scale bar: 250ms). **b.** Schematic of a horizontal section showing the center of recording site (Left). Representative horizontal section showing post-mortem histology illustrating probe location(Right). Blue is a glial marker (CD11b) and green is NeuN. Scale bar: 100um. **c.** Scatter plot showing the classification of isolated units based on their waveform characteristics to putative regular spiking (RS) and fast-spiking (FS) cells. **d**,**e.** Example cells from RS (top) and FS (bottom) showing heterogeneous selectivity for ipsilateral and contralateral choices at different time points within trials.

We classified recorded units based on their waveform shape to belong to either putative regular spiking (RS) or fast-spiking (FS) neurons (Fig. 5c). Examples of spiking activity of each cell type is shown in Fig 5d. As expected in prefrontal cortex, activity profiles were quite heterogeneous among individual units. A significant proportion of cells (45%) were modulated beyond their baseline firing rate (random shuffle test comparing 30ms bins to pre-fixation period) but briefly and at different timepoints, spanning the entire duration of the trial (Fig. 6a). Although we did not find evidence of persistently active single units during the delay period, RS cells as a population were persistently active during this epoch (Fig. 6b). FS cells, in contrast, did not show persistent activity during the delay period and became disproportionately active during the choice execution.

**Fig. 6.**
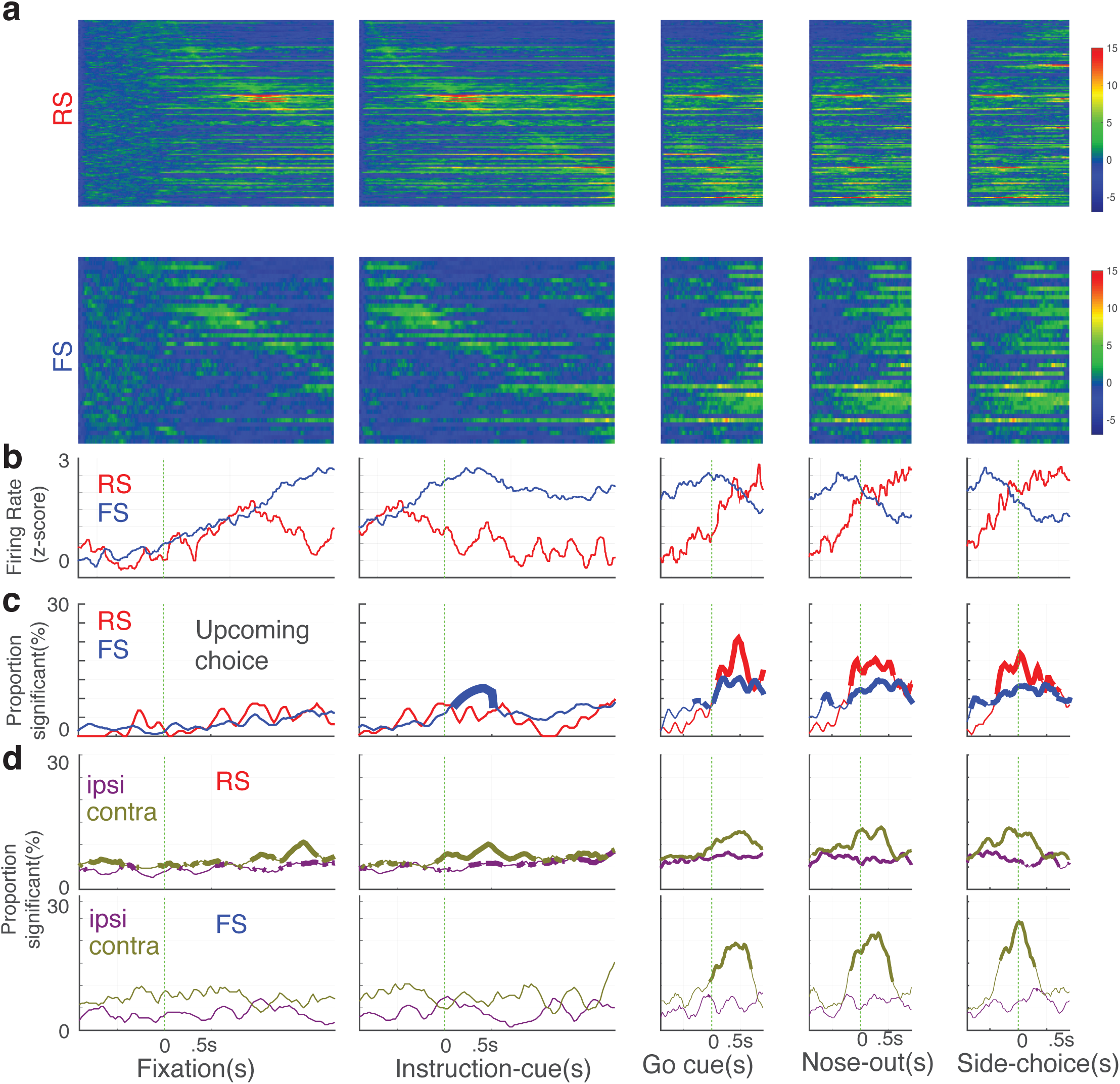
RS cells selectively encode the contralateral choice during instruction. **a.** Population response of mPFC cells during different task epochs (z-score). Most cells are selective only during a brief epoch in a given trial. **b.** Average population response of RS and FS cells at different time points. RS cells collectively show persistent activity during the delay period. However, FS cells ramp up their activity after the Go cue and during choice execution. **c.** The proportion of units selective for choice at different time points. Both RS and FS cells are significantly selective (p<.01, binomial test, n=155 for RS cells and n=44 for FS cells) after the go cue. However, only RS cells are selective during the instruction cue. Thick lines demonstrate time points with significantly proportion of cells. **d.** Proportion of individual cells selective for ipsilateral and contralateral trials at different time points. RS cells significantly encode contralateral trials during the instruction cue.

Both RS and FS population encoded information regarding upcoming choice after the Go cue (i.e., during choice execution). However, only RS cells significantly encoded the upcoming choice during the instruction period (Fig. 6c). Also, both populations encoded the outcome (rewarded/unrewarded) after the trial outcome was revealed to the animal. At no point during the trial (or inter-trial-interval) a significant population of RS or FS cells carried information about the previous trial choice or outcome (p>0.05, binomial test). Finally, RS (but not FS) cells encoded information about contralateral choices during the delay period (Fig. 6d).

Given the heterogeneous nature of response encoding by individual cells at different timepoints during each trial, we then turned to population-level analysis to investigate PFC dynamics. Reduced dimension trajectories were constructed for contralateral and ipsilateral trials using Principal Component Analysis (PCA) projections. Dimensionality reduction was performed on smoothed firing rate using a gaussian kernel with a 25ms width. During fixation (and before the instruction cue), there was small variability in the population response and the trajectories for contra and ipsi trials were almost indistinguishable from one another (Fig. 7a). At the instruction cue onset, however, population trajectory deviated significantly from baseline (p<0.01, shuffle test, n=1000), but only for contralateral choice trials, suggesting a decision is made about an upcoming choice. By the time go cue arrives, distinct trajectories have evolved both for ipsilateral and contralateral trials. These trajectories remained separate until the outcome of the trial was revealed at which point the population activity returned to their initial state.

**Fig. 7.**
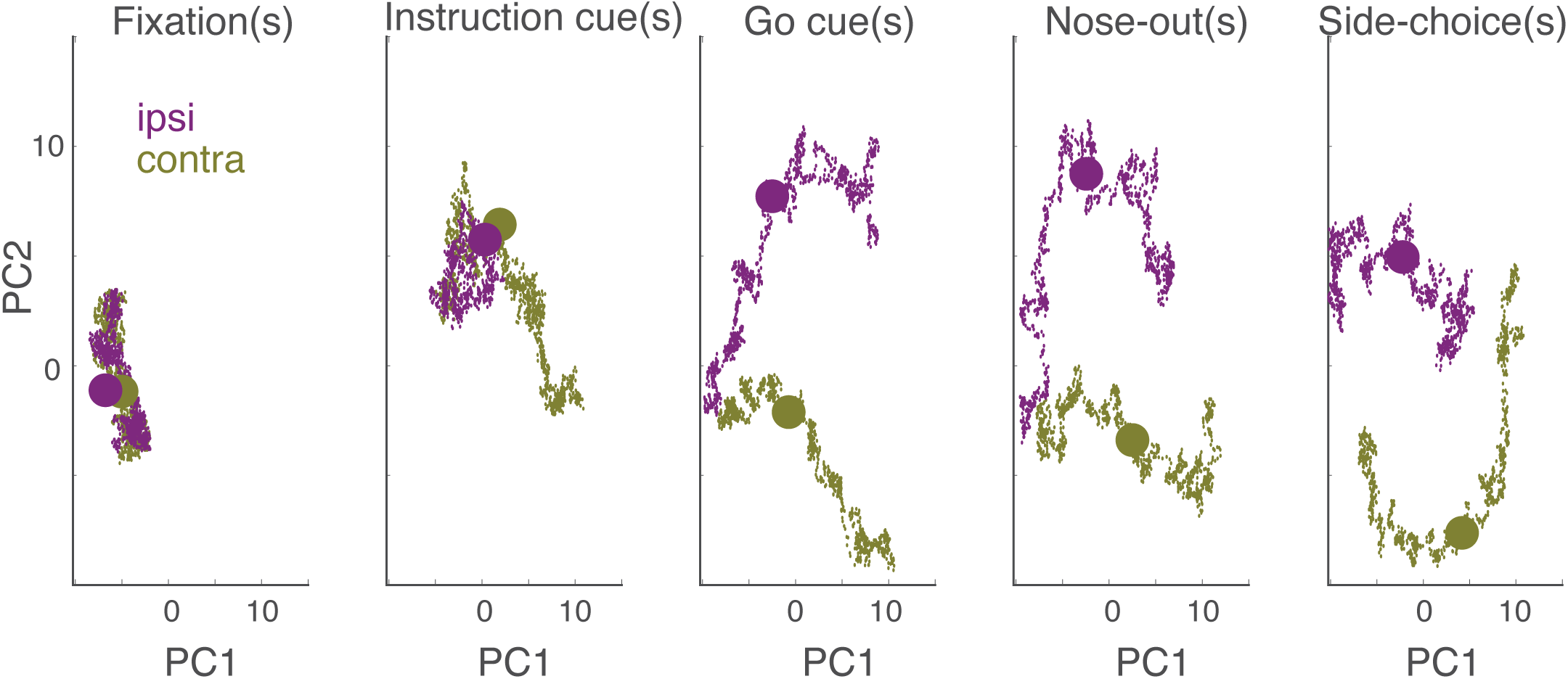
Population dynamic state space underlying choice representation in the prelimbic cortex. Reduced dimension representation of population activity around key trial events visualized in a two-dimensional principal component space. Each dot represents the two-dimensional projection of the population dynamics, obtained from smoothed firing rate using a Gaussian kernel of 25ms width sampled at 1ms intervals. The trajectories are plotted for .5s before and after each trial event (denoted by a large circle for contralateral and ipsilateral trials). During fixation, population dynamics show small deviation from baseline. The introduction of the instruction cue, however, triggers a deviation from an initial state but only for contralateral trials. By the time the subject plans the movement and awaits the go cue, population trajectories become completely separated and distinct from baseline. Trajectories continue to be distinct until the animal fully commits to a choice.

## DISCUSSION

Prefrontal cortex plays an important role in a wide range of cognitive behaviors. Here, we sought to investigate its role in sensory-guided orienting behavior in rats. We trained rats to perform a delayed two-alternative forced-choice task. Using two independent manipulation methods and multi-electrode recordings in the prelimbic cortex, our data reveals a previously uncharacterized role of mPFC in top-down control of decision making during orienting behavior. We first demonstrated that unilateral suppression of mPFC activity during whole trial intervals impairs contralateral choices, consistent with previous reports (Erlich et al., 2011). We further found that precise photo-suppression during different epochs of the task have dichotomous effects on choice. Specifically, activity suppression during the instruction period only affected contralateral choices while delay period suppression equally affected both contralateral and ipsilateral choices. We explained this apparent paradox by recording from mPFC cells and demonstrating that both single unit and population dynamics deviate from baseline only for contralateral trials during the instruction cue. This role is primarily carried out by putative RS pyramidal cells. RS cells constitute a primarily excitatory neuron cell type that were targeted by our ArchT-mediated suppression protocol design using the CaMKIIa promoter. Based on this demonstration, we propose that mPFC plays an asymmetrical role in orienting behavior during the stage in which extraction of salient features from sensory input takes place to guide decisions about future orienting movement. These results are in contrast with previous reports investigating the role of other subregions of PFC in orienting behavior. In particular, whereas prior work targeted the so called “frontal orienting field (FOF)” (AP:2.0, ML:1.3, DV:-0.8mm), we have targeted the prelimbic area (AP:2.6-4.0, ML:0.8:DV:-3.00mm), which is considered an upstream target of FOF neurons. In addition, while FOF has been labeled a premotor area dedicated to preparation of orientating movements, the prelimbic area is thought to be more involved in sensorimotor integration, reasoning and rule retrieval (Brasted et al., 2000; Duan et al., 2015; Euston et al., 2012; Hardung et al., 2017; Kamigaki and Dan, 2017; Kim et al., 2016; Sul et al., 2010). As such, the distinct patterns of activity in the prelimbic cortex during the delay period compared to the instruction period are likely serving different roles in mediating choice behavior.

We observed persistent activity during the delay period only at the population level but not at individual cells(Stokes, 2015; Stokes et al., 2017). Prior modeling studies have emphasized the role of persistent activity of prefrontal neurons in the representation of decisions in working memory (Bernacchia et al., 2011; Daie et al., 2015; Soltani et al., 2006). These studies were based on earlier neurophysiological observations of memory cells in primates (Funahashi, 2006; Funahashi et al., 1989; Fuster et al., 1971) in which a balance between excitation and inhibition was proposed to maintain such representations (Lim and Goldman, 2013; Murray et al., 2017; Wang et al., 2004). A recent modeling study have suggested that depending on the task requirements, mPFC cells may or may not demonstrate persistent activity (Masse et al., 2018). As such, it remains a topic of debate whether persistent activity is a characteristic of mPFC cells or a consequence of biased observations (Arnsten et al., 2010; Constantinidis et al., 2018; Lundqvist et al., 2018; Stokes, 2015; Stokes et al., 2017).

In our study, however, only putative RS pyramidal cells demonstrated persistent activity. While individual RS cells were choice selective as early as the instruction cue onset, they demonstrated sustained activity throughout the delay period, albeit in the form of short, transiently active clusters of cells. FS cells, on the other hand, were only recruited after the go cue during choice execution. Anatomical studies suggest that the majority of FS cells are Parvalbumin (PV)-expressing interneurons targeting the somatic compartments of excitatory projection cell bodies (Kvitsiani et al., 2013; Pi et al., 2013; Pinto and Dan, 2015). This connectivity would enable them to rapidly suppress projection cell activities and thus exert significant control of the output of mPFC circuits. This might explain the activity pattern of FS cells observed around the choice execution: suppressing an unwanted element of a potential motor plan in order to bias the motor system towards executing the appropriate orienting movement. This role would be consistent with projection pathways from mPFC (Chatham et al., 2014) as well as other parts of cortex (e.g. posterior parietal cortex PPC(Hwang et al., 2019; Lyamzin and Benucci, 2019) to the basal ganglia that facilitate desired movements and suppress undesired movements. Given that suppression of excitatory neural activity in mPFC during the motor planning stage affected both contralateral and ipsilateral movements, our data suggest that the response selectivity of RS and FS neurons contribute to this parallel but coordinated elements of the motor plan.

We limited our recordings to layer 5 of the dorsal prelimbic cortex since it comprises most projection cells that send efferents downstream to other cortical (particularly PPC (Denardo et al., 2015) and subcortical regions (particularly striatal regions) and thus may be directly responsible for top-down control signals that affect orienting decisions. As such, it is likely that pharmacological inactivation effects might have extended outside of this layer. It remains to be examined in future studies whether projection targets of these pyramidal cells can explain their selectivity for ipsilateral and contralateral choices. For example, it is quite possible that intratelencephalic (IT) and pyramidal tract (PT) neurons disproportionately represent contralateral and ipsilateral choices, respectively(Shepherd, 2013). Using optogenetic identification combined with retrograde transport may help resolve such dichotomies (Anikeeva et al., 2011; Kvitsiani et al., 2013; Tervo et al., 2016).

It would also be of interest to compare the population dynamics in layer 2/3 of dorsal prelimbic cortex. While we did not find representations of ipsilateral choice during the instruction cue in layer 5 responses, layer 2/3 that directly receives more sensory afferents compared to layer 5 may carry such representations. Layer 5 and layer 2/3 also have distinct interneuronal cytoarchitecture, with layer 2/3 containing disproportionately more VIP and SOM interneurons compared to dense presence of PV interneurons in layer 5 (Gabbott et al., 1997). In contrast to PV cells, VIP and SOM cells target distal dendrites (Kepecs and Fishell, 2014; Pi et al., 2013). Given these differences, we predict that layer 2/3 interneurons help sculpt sensorimotor transformation whereas layer 5 FS cells (mostly PV) inhibit the outputs of these circuits and thus are only active during choice execution, as we demonstrated in the present study.

Our unit recording data confirmed the heterogeneous activity patterns reported by many previous studies, suggestive of network dynamics encoding (Rigotti et al., 2013). However, it is possible that this heterogeneous response could be explained by the phenotype and projection patterns of these mPFC cells (Hirokawa et al., 2019) as discussed above. Further experiments with cell-type and projection specific targeting of individual cells are required to test the origin of this heterogeneity in network patterns in the context of similar task design.

## METHODS

### Two-alternative forced-choice task

We used a two-alternative forced-choice task design similar to previously published studies (Brunton et al., 2013; Gage et al., 2010). Adult Sprague-Dawley rats (Charles-River) were food deprived to their 85% ad libitum weight and placed in acoustically isolated training boxes (Coulbourn instruments, PA). The box comprised an operant conditioning chamber with three nosepoke holes (a fixation hole in the center and two target holes on the sides) and a reward delivery trough on the opposite side of the nose poke holes. A speaker was centrally placed on the reward trough side and used to deliver auditory cues to the subject. Subjects were required to place their nose inside the fixation hole for 1s waiting for an instruction cue. Instruction cue was a brief (.5s) single tone delivered at 60 dB. A low pitch tone (5 KHz) indicated an instruction to choose the right nose poke hole, whereas a 1.5 octave higher-tone (14.2 KHz) indicated an instruction to choose the left nose poke hole. The instruction cue was followed by a delay period of variable length (chosen pseudo-randomly from a uniform distribution between 1-1.5s), which was followed by a Go cue-a white noise stimulus at 60dB. If the subject retracted from the fixation hole before the Go cue, the trial was terminated and considered a premature retraction. Visits to the instructed targets were rewarded immediately by delivering a 45mg food pellet, while incorrect visits were punished by extending the inter-trial-interval from 7s for successful trials to 12s for failed trials with no reward pellet delivery. Stimulus generation, behavioral state control and reward delivery were all conducted using the Coulbourn habitest system (Coulbourn Instruments, PA) and the data was logged in real-time on a desktop computer. All behavioral events were sampled at 1KHz and subsequent analysis was performed using custom-developed MATLAB routines.

### Microelectrode array implant surgery

Once subjects became proficient in the task and maintained a steady performance score above 85% correct, they were removed from the food deprivation protocol for a week prior to the surgery day. Subjects were anesthetized using inhalation of 2% Isofluorane in an induction chamber. Body temperature and vital signs were monitored throughout the surgery. A craniotomy was performed on top of the medial prefrontal cortex (+2-5mm AP, 0.5-1.5mm ML) to expose the brain surface. A 32 channel microwire array (Tucker-Davis Technologies) was slowly advanced into the cortex. Extracellular potentials were recorded during the penetration procedure with respect to cerebellar ground and reference screws touching the dura matter posterior to the lambda point. Local Field Potentials (LFPs) and multiple single-unit activity were monitored until the probe reached the target depth. The craniotomy was covered with GelFoam (Pfizer, NY) and then the probe was anchored to the skull by applying dental cement (C&B metabond, Parkell, NY) using 6 skull screws surrounding the probe. Subjects were then given one full week to recover from surgery before being subjected again to the food deprivation protocol.

### Reversible inactivation

For the muscimol inactivation studies, cannulas were placed on top of the prelimbic cortex (AP:3.00, ML:0.8, DV:-3.2) anchored to the skull using stainless steel screws and dental cement. Rats were allowed two weeks to recover and rehabilitate in the task. Prior to the infusions, rats were lightly anesthetized using isoflurane. The dummy cannula was removed and replaced on one side with an injector (Hamilton Company, Nevada, USA) the end of which extended 0.5 mm from the tip of the guide cannula. GABAa receptor agonist muscimol (Sigma, Missouri, USA) was infused into the mPFC to reversibly inactivate this region. Muscimol solution (1 ug/uL in artificial cerebrospinal fluid, 1.0 μL) was infused unilaterally into one side of the cannula. The day prior to and the day following the muscimol infusion, an ACSF (1.0 uL) solution was unilaterally infused into the same side as a control. Infusions were performed at the slow rate of .1625 uL/min, and the infusion needle remained in place for 10 minutes following the injection. Rats performed the task 90 minutes after infusion for 90 minutes. One week after the first ACSF infusion, the infusion procedure was repeated on the other side of the brain.

### Optogenetic suppression

Trained rats with performance scores above 85% were selected for the optogenetic experiment. Standard surgical procedures were followed for intracranial injection. Glass pipettes were pulled to OD of ∼40um with a long taper and penetrated in the brain via bare holes made on top of the prelimbic cortex, bilaterally. Each injection delivered 1uL of AAV5-CaMKIIa-ArchT-eYFP (Han et al., 2011) at the slow rate of 1uL/10 min at the same coordinates as muscimol injections. Ferrule attached optical fibers (NA=.48, 200um in diameter, Doric Lenses, ON) were then implanted stereotaxically, sitting on top of the infected area. Ferrules were then connected to diode pump solid state lasers (either 446nm or 520nm) delivering laser pulses at 10mW.

### Electrophysiological recordings

Data acquisition was performed using a Tucker-Davis RZ2 through a high impedance headstage and an automated commutator to permit unrestricted movement in the behavioral box. Wide-band signals were recorded at the rate of 25KHz/channel, amplified and digitized to 16-bits. LFPs were separated from multi-unit activity using a Symlet wavelet filtering algorithm. Once filtered, the noise root mean square value was calculated from the raw data and samples surpassing a threshold 4 times this noise floor was used as an indication of action potential occurrence. Extracted waveforms were then sorted using a custom-developed software – referred to as ‘EZsort’-in MATLAB.

### Histology

Subjects animals were deeply anesthetized and transcardially perfused with 4% paraformaldehyde prepared in 0.1M phosphate-buffered saline (PBS). Brains were explanted and post-fixed in 4% paraformaldehyde overnight at 4C. Tissue was sectioned on a vibrating microtome at a 50-micron thickness, where coronal sections were used to identify ArchT/GFP-expressing neurons, and sections transverse to the probe shank were used to validate the implant location. For optogenetic experiments, sections were counterstained with Hoechst (1 ug/mL, 10 minutes), rinsed and mounted with ProLong Gold Anti-fade mounting medium (Invitrogen, Carlsbad, CA). For probe location, sections were rinsed, blocked with 10% normal goat serum, and incubated overnight at 4C in a solution of mouse-anti-CD11b (abcam) to identify the microglial sheath (1:100 dilution in 0.3% triton, 5% normal goat serum in PBS)(Ward et al., 2009). Sections were subsequently labeled with goat-anti-mouse 488 (Invitrogen, Carlsbad, CA)(1:200 dilution in 0.3% triton, 5% normal goat serum in PBS), counterstained with Hoechst (1 ug/mL, 10 minutes), rinsed and mounted with ProLong Gold Anti-fade mounting medium. Results were imaged on an Olympus fluoview confocal microscope. For optogenetic studies, an observer blinded to the experimental condition evaluated the expression level. Only those subjects with observable expression were included in the study.

## ACKNOWLEDGEMENTS

The authors thank Dylan Miller for technical assistance in the pharmacological inactivation experiment and Erin K. Purcell for assistance in viral injections and histology. This work was supported by the National Institute on Drug Abuse, the National Institutes Health, through NS93909 and NS062031 grants.

## AUTHOR CONTRIBUTIONS

A.M. and K.G.O. designed the experiments. A.M. perfomred the experiments and analyzed the data. K.G.O. supervised the study and A.M. And K.G.O. wrote the manuscript.

## COMPETING INTERESTS

The authors decalre no competing interests.

**Correspondence and requests for materials** should be addressed to A.M.

## Notes

### Competing Interest Statement

The authors have declared no competing interest.

## REFERENCES

Anikeeva, P., Andalman, A.S., Witten, I.B.B., Warden, M., Goshen, I., Grosenick, L., Gunaydin, L.A., Frank, L.M., and Deisseroth, K. (2011). Optetrode: a multichannel readout for optogenetic control in freely moving mice. Nat. Neurosci. 15, 163–170.

Arnsten, A.F.T., Paspalas, C.D., Gamo, N.J., Yang, Y., and Wang, M. (2010). Dynamic network connectivity: A new form of neuroplasticity. Trends Cogn. Sci. 14, 365–375.

Baker, P.M., and Ragozzino, M.E. (2014). Contralateral disconnection of the rat prelimbic cortex and dorsomedial striatum impairs cue-guided behavioral switching. Learn. Mem. 21, 368–379.

Bernacchia, A., Seo, H., Lee, D., and Wang, X. (2011). A reservoir of time constants for memory traces in cortical neurons. Nat. Neurosci. 14, 366–372.

Brasted, P.J., Dunnett, S.B., and Robbins, T.W. (2000). Unilateral lesions of the medial agranular cortex impair responding on a lateralised reaction time task. Behav. Brain Res. 111, 139–151.

Brunton, B.W., Botvinick, M.M., and Brody, C.D. (2013). Rats and humans can optimally accumulate evidence for decision-making. Science 340, 95–98.

Chatham, C.H., Frank, M., and Badre, D. (2014). Corticostriatal Output Gating during Selection from Working Memory. Neuron 81, 930–942.

Constantinidis, C., Funahashi, S., Lee, D., Murray, J.D., Qi, X.L., Wang, M., and Arnsten, A.F.T. (2018). Persistent spiking activity underlies working memory. J. Neurosci. 38, 7020–7028.

Daie, K., Goldman, M.S., and Aksay, E.R.F. (2015). Spatial Patterns of Persistent Neural Activity Vary with the Behavioral Context of Short-Term Memory. Neuron 85, 847–860.

Denardo, L.A., Berns, D.S., Deloach, K., and Luo, L. (2015). Connectivity of mouse somatosensory and prefrontal cortex examined with trans-synaptic tracing. Nat. Neurosci. 18, 1687–1697.

Donahue, C.H., and Lee, D. (2015). Dynamic routing of task-relevant signals for decision making in dorsolateral prefrontal cortex. Nat. Neurosci. 18.

Duan, C.A., Erlich, J.C., and Brody, C.D. (2015). Requirement of Prefrontal and Midbrain Regions for Rapid Executive Control of Behavior in the Rat. Neuron 86, 1491–1503.

Erlich, J.C., Bialek, M., and Brody, C.D. (2011). A cortical substrate for memory-guided orienting in the rat. Neuron 72, 330–343.

Euston, D.R., Gruber, A.J., and McNaughton, B.L. (2012). The Role of Medial Prefrontal Cortex in Memory and Decision Making. Neuron 76, 1057–1070.

Fujisawa, S., Amarasingham, A., Harrison, M.T., and Buzsáki, G. (2008). Behavior-dependent short-term assembly dynamics in the medial prefrontal cortex. Nat. Neurosci. 11, 823–833.

Funahashi, S. (2006). Prefrontal cortex and working memory processes. Neuroscience 139, 251–261.

Funahashi, S., Bruce, C.J., and Goldman-Rakic, P.S. (1989). Mnemonic coding of visual space in the monkey’s dorsolateral prefrontal cortex. J. Neurophysiol. 61, 331–349.

Fuster, J.M., Alexander, G.E., and others (1971). Neuron activity related to short-term memory. Science (80-.). 173, 652–654.

Gabbott, P.L.A., Dickie, B.G.M., Vaid, R.R., Headlam, A.J.N., and Bacon, S.J. (1997). Local-Circuit Neurones in the Medial Prefrontal Cortex (Areas 25, 32 and 24b) in the Rat: Morphology and Quantitative Distribution Indexing terms: calcium-binding proteins; GABA; NADPH diaphorase; cortical modules; limbic system. J. Comp. Neurol 377, 465–499.

Gabbott, P.L.A., Warner, T.A., Jays, P.R.L., Salway, P., and Busby, S.J. (2005). Prefrontal cortex in the rat: projections to subcortical autonomic, motor, and limbic centers. J. Comp. Neurol. 492, 145–177.

Gage, G.J., Stoetzner, C.R., Wiltschko, A.B., and Berke, J.D. (2010). Selective activation of striatal fast-spiking interneurons during choice execution. Neuron 67, 466–479.

Gisquet-Verrier, P., and Delatour, B. (2006). The role of the rat prelimbic/infralimbic cortex in working memory: not involved in the short-term maintenance but in monitoring and processing functions. Neuroscience 141, 585–596.

Greene, J.D., Sommerville, R.B., Nystrom, L.E., Darley, J.M., and Cohen, J.Y. (2001). An fMRI investigation of emotional engagement in moral judgment. Science 293, 2105–2108.

Han, X., Chow, B.Y., Zhou, H., Klapoetke, N.C., Chuong, A., Rajimehr, R., Yang, A., Baratta, M. V, Winkle, J., Desimone, R., et al. (2011). A high-light sensitivity optical neural silencer: development and application to optogenetic control of non-human primate cortex. Front. Syst. Neurosci. 5, 18.

Hardung, S., Epple, R., Jäckel, Z., Eriksson, D., Uran, C., Senn, V., Gibor, L., Yizhar, O., and Diester, I. (2017). A Functional Gradient in the Rodent Prefrontal Cortex Supports Behavioral Inhibition. Curr. Biol. 27, 549–555.

Hirokawa, J., Vaughan, A., Masset, P., Ott, T., and Kepecs, A. (2019). Frontal cortex neuron types categorically encode single decision variables. Nature 576, doi.org/10.1038/s41586-019-1816-9.

Hwang, E.J., Link, T.D., Hu, Y.Y., Neil, K.O., Hwang, E.J., Link, T.D., Hu, Y.Y., Lu, S., Wang, E.H., and Lilascharoen, V. (2019). Corticostriatal Flow of Action Selection Bias Article Corticostriatal Flow of Action Selection Bias. Neuron 1–15.

Kamigaki, T., and Dan, Y. (2017). Delay activity of specific prefrontal interneuron subtypes modulates memory-guided behavior. Nat. Neurosci. 20, 854–863.

Karlsson, M.P., Tervo, D.G.R., and Karpova, A.Y. (2012). Network Resets in Medial Prefrontal Cortex Mark the Onset of Behavioral Uncertainty. Science (80-.). 338, 135–139.

Kepecs, A., and Fishell, G. (2014). Interneuron cell types are fit to function. Nature 505, 318–326.

Kim, D., Jeong, H., Lee, J., Ghim, J., Her, E.S., Lee, S., Kim, D., Jeong, H., Lee, J., Ghim, J., et al. (2016). Distinct Roles of Parvalbumin- and Somatostatin-Expressing Interneurons in Working Memory. Neuron 1–14.

Kopec, C.D., Erlich, J.C., Brunton, B.W., Deisseroth, K., and Brody, C.D. (2015). Cortical and Subcortical Contributions to Short-Term Memory for Orienting Movements Article Cortical and Subcortical Contributions to Short-Term Memory for Orienting Movements. Neuron 1–11.

Kvitsiani, D., Ranade, S., Hangya, B., Taniguchi, H., Huang, J.Z., and Kepecs, A. (2013). Distinct behavioural and network correlates of two interneuron types in prefrontal cortex. Nature 498, 363–366.

Lim, S., and Goldman, M.S. (2013). Balanced cortical microcircuitry for maintaining information in working memory. Nat. Neurosci. 1–11.

Lundqvist, M., Herman, P., and Miller, E.K. (2018). Working memory: Delay activity, yes! persistent activity? maybe not. J. Neurosci. 38, 7013–7019.

Lyamzin, D., and Benucci, A. (2019). The mouse posterior parietal cortex: Anatomy and functions. Neurosci. Res. 140, 14–22.

Masse, N.Y., Yang, G.R., Song, H.F., Wang, X.-J., and Freedman, D.J. (2018). Circuit mechanisms for the maintenance and manipulation of information in working memory. Nat. Neurosci.

Miller, E.K., and Cohen, J.Y. (2001). An integrative theory of prefrontal cortex function. Annu. Rev. Neurosci. 24, 167–202.

Mohebi, A., and Oweiss, K.G. (2014). A fully automated rodent conditioning protocol for sensorimotor integration and cognitive control experiments. J. Vis. Exp. 86.

Murray, J.D., Bernacchia, A., Roy, N.A., Constantinidis, C., Romo, R., and Wang, X.J. (2017). Stable population coding for working memory coexists with heterogeneous neural dynamics in prefrontal cortex. Proc. Natl. Acad. Sci. U. S. A. 114, 394–399.

Narayanan, N.S., and Laubach, M. (2006). Top-down control of motor cortex ensembles by dorsomedial prefrontal cortex. Neuron 52, 921–931.

Pi, H.J., Hangya, B., Kvitsiani, D., Sanders, J.I., Huang, Z.J., and Kepecs, A. (2013). Cortical interneurons that specialize in disinhibitory control. Nature 503, 521–524.

Pinto, L., and Dan, Y. (2015). Cell-Type-Specific Activity in Prefrontal Cortex during Goal-Directed Behavior. Neuron 87, 437–450.

Powell, N.J., and Redish, A.D. (2016). Representational changes of latent strategies in rat medial prefrontal cortex precede changes in behaviour. Nat. Commun. 7, 12830.

Quintana, J., and Fuster, J.M. (1999). From perception to action: temporal integrative functions of prefrontal and parietal neurons. Cereb. Cortex 9, 213–221.

Rich, E.L., and Shapiro, M.L. (2009). Rat Prefrontal Cortical Neurons Selectively Code Strategy Switches. J. Neurosci. 29, 7208–7219.

Rigotti, M., Barak, O., Warden, M.R., Wang, X., Daw, N.D., Miller, E.K., and Fusi, S. (2013). The importance of mixed selectivity in complex cognitive tasks. Nature.

Romo, R., and de Lafuente, V. (2013). Conversion of sensory signals into perceptual decisions. Prog. Neurobiol. 103, 41–75.

Shepherd, G.M.G. (2013). Corticostriatal connectivity and its role in disease. Nat. Rev. Neurosci. 14, 278–291.

Soltani, A., Lee, D., and Wang, X. (2006). Neural mechanism for stochastic behaviour during a competitive game. Neural Netw. 19, 1075–1090.

Stokes, M.G. (2015). “Activity-silent” working memory in prefrontal cortex: A dynamic coding framework. Trends Cogn. Sci. 19, 394–405.

Stokes, M.G., Buschman, T.J., and Miller, E.K. (2017). Dynamic Coding for Flexible Cognitive Control. 221–241.

Sul, J.H., Kim, H., Huh, N., Lee, D., and Jung, M.W. (2010). Distinct roles of rodent orbitofrontal and medial prefrontal cortex in decision making. Neuron 66, 449–460.

Szczepanski, S.M., and Knight, R.T. (2014). Insights into Human Behavior from Lesions to the Prefrontal Cortex. Neuron 83, 1002–1018.

Tervo, D.G.R., Hwang, B.Y., Viswanathan, S., Gaj, T., Lavzin, M., Ritola, K.D., Lindo, S., Michael, S., Kuleshova, E., Ojala, D., et al. (2016). A Designer AAV Variant Permits Efficient Retrograde Access to Projection Neurons. Neuron 92, 372–382.

Wang, X.J., Tegnér, J., Constantinidis, C., and Goldman-Rakic, P.S. (2004). Division of labor among distinct subtypes of inhibitory neurons in a cortical microcircuit of working memory. Proc. Natl. Acad. Sci. U. S. A. 101, 1368–1373.

Willcocks, A.L., and McNally, G.P. (2013). The role of medial prefrontal cortex in extinction and reinstatement of alcohol-seeking in rats. Eur. J. Neurosci. 37, 259–268.

